# Raman Needle Arthroscopy for *In Vivo* Molecular Assessment of Cartilage

**DOI:** 10.1101/2021.06.15.448529

**Authors:** Kimberly Kroupa, Man I Wu, Juncheng Zhang, Magnus Jensen, Wei Wong, Julie B. Engiles, Mark W. Grinstaff, Brian D. Snyder, Mads S. Bergholt, Michael B. Albro

**Author notes:** denotes equal contribution (Kimberly Kroupa, Man I Wu, Juncheng Zhang).

## Abstract

The development of treatments for osteoarthritis (OA) is burdened by the lack of standardized biomarkers of cartilage health that can be applied in clinical trials. We present a novel arthroscopic Raman probe that can “optically biopsy” cartilage and quantify key ECM biomarkers for determining cartilage composition, structure, and material properties in health and disease. Technological and analytical innovations to optimize Raman analysis include: 1) multivariate decomposition of cartilage Raman spectra into ECM-constituent-specific biomarkers (glycosaminoglycan [GAG], collagen [COL], water [H_2_O] scores), and 2) multiplexed polarized Raman spectroscopy to quantify superficial zone collagen anisotropy via a PLS-DA-derived Raman collagen alignment factor (RCAF). Raman measurements were performed on a series of *ex vivo* cartilage models: 1) chemically GAG-depleted bovine cartilage explants (n=40), 2) mechanically abraded bovine cartilage explants (n=30), 3) aging human cartilage explants (n=14), and 4) anatomical-site-varied ovine osteochondral explants (n=6). Derived Raman GAG score biomarkers predicted 95%, 66%, and 96% of the variation in GAG content of GAG-depleted bovine explants, human explants, and ovine explants, respectively (p<0.001). RCAF values were significantly different for explants with abrasion-induced superficial zone collagen loss (p<0.001). The multivariate linear regression of Raman-derived ECM biomarkers (GAG and H_2_O scores) predicted 94% of the variation in elastic modulus of ovine explants (p<0.001). Finally, we demonstrated the first *in vivo* Raman arthroscopy assessment of an ovine femoral condyle through intraarticular entry into the synovial capsule. This work advances Raman arthroscopy towards a transformative low cost, minimally invasive diagnostic platform for objective monitoring of treatment outcomes from emerging OA therapies.

## Introduction

Osteoarthritis (OA) is a chronic, debilitating condition, characterized by the progressive degradation of articular cartilage. It is the most widespread cause of disability in adults; over 20% of the US adult population (>50 million individuals) is afflicted with the disease, with this number predicted to rise sharply over coming decades ^1,2^. Currently, no FDA approved therapies exist that reliably mitigate the degradation of cartilage tissue properties induced by OA.

The structure and composition of articular cartilage is optimized for its mechanical performance. It is comprised of a type-II collagen (COL) fibril network that affords structure and tensile strength, complemented by a negatively charged sulfated glycosaminoglycan (GAG) matrix that provides compressive properties and retains interstitial water ^3^. More than 90% of applied joint load is supported by pressurization of entrapped water (interstitial fluid load support), yielding the tissue’s characteristic low frictional properties ^4^. Additionally, cartilage is inhomogeneous and structurally anisotropic, where the collagen configuration varies with depth, segregated into zones optimized for mechanical performance: *superficial zone (SZ)* - comprised of collagen fibers arranged parallel to the articular surface, *middle zone (MZ)* - mix-aligned collagen, and *deep zone (DZ)* - perpendicular arranged collagen. During OA, the GAG and collagen constituents of the cartilage matrix become depleted, leading to compromised mechanical tissue function. Early on, GAG is depleted from the SZ ^5,6^ with concomitant loss of superficial collagen fiber organization and alignment ^7,8^. Loss of GAG and COL alignment reduces fluid load support, transferring loads to the collagen matrix, and leading to cartilage erosion through the MZ and DZ. Later stages of OA progression are characterized by further GAG depletion and surface delamination, culminating in significant cartilage volume loss until bone-on-bone contact is reached ^9^.

Numerous potential strategies to mitigate, protect, or regenerate the material properties of articular cartilage are emerging, including 1) modified joint kinetics (e.g., weight loss ^10^, physical therapy ^11^), 2) disease modifying OA drugs ^12^, 3) viscosupplements ^13^, and 4) surgically assisted tissue regeneration (microfracture, chondrocyte implantation, stem cell therapies) ^14^. However, the ability to assess the efficacy of OA treatments that preserve and/or regenerate cartilage tissue structure and function is burdened by a lack of standardized biomarkers that can be applied in preclinical and clinical trials. Notably, clinical trials evaluating the efficacy of therapies rely on patient pain and mobility (WOMAC) scores that can be contaminated by placebo effect and often correlate poorly to objective metrics of cartilage composition and structure. Image-based assessments (radiographs and MRI) are biased towards late stage OA pathoanatomy (cartilage volume loss, bone marrow edema, cysts, and osteophytes). MRI measurements of cartilage composition and structure may be affected by lack of consistency, SNR, and insufficient image resolution.

Raman spectroscopy is predicated on the inelastic scattering of photons. When monochromic laser light induces a change in molecular polarizability during vibrations, a small proportion of the incident photons (~1 in 10^8^) are scattered with a change in wavelength ^15^. The Raman scattered light reflects the vibrational modes of constituent molecules; the absorbed energy corresponds to specific Raman active vibrational modes that define a molecule’s “fingerprint”. Thus, Raman spectra carry information about individual molecular vibrational bonds that correspond to specific biochemical building blocks (amides, sulfates, carboxylic acids, hydroxyls) of key cartilage constituents (GAG, COL, H_2_O). The premise of this study is that Raman spectroscopy can “optically biopsy” cartilage and quantify the relative contribution of key constituents that serve as biomarkers for determining cartilage composition, structure, and material properties in health and disease. Prior work has demonstrated that Raman spectra of cartilage exhibit statistical changes in response to mechanical damage ^16-18^ and OA ^19-22^. However, the implementation of Raman spectroscopy as a diagnostic tool for cartilage health has been impeded by a lack of clinically compatible intra-articular Raman probes for *in vivo* diagnostics and an inability to extract specific and quantitative biochemical and structural metrics related to cartilage health.

In this work we develop a clinically-compatible Raman needle arthroscopy platform for *in vivo* molecular assessment of cartilage. A thin, fiber-optic Raman spectroscopic probe, which can be directed intra-articularly via a hypodermic-needle-cannula, was developed to “optically biopsy” cartilage at specific anatomic sites under image guidance. Technological and analytical innovations to optimize Raman spectral analysis include: 1) decomposing composite Raman cartilage spectra and isolating the relative contribution of specific cartilage constituents, and 2) using multiplexed polarized Raman spectroscopy to quantify SZ collagen anisotropy. We first establish the potential of our Raman arthroscopic probe to measure composition, structure, and material properties of cartilage that undergo changes during OA through parametric analysis of a series of *ex vivo* model systems: 1) Raman quantification of GAG content in enzymatically depleted bovine cartilage and aging human cartilage explants, 2) Raman quantification of zonal collagen network organization in mechanically abraded bovine cartilage explants, and 3) Raman measurements of bovine cartilage thickness. Subsequently, we assess the ability of Raman biomarkers to predict the composition, thickness, and mechanical properties of cartilage explants from different anatomical sites of an ovine knee joint. We finally perform an *in vivo* Raman arthroscopic assessment of the composition of an ovine femoral condyle, demonstrating the clinical feasibility of Raman OA diagnostics.

## Methods

### Raman needle arthroscopic probe instrumentation

A custom polarized Raman needle arthroscopic probe was developed for *in vivo* OA diagnostics by intra-articular entry through a hypodermic needle (Fig.1a-c) ^23^ (described in *Supplementary Material*).

**Fig 1.**
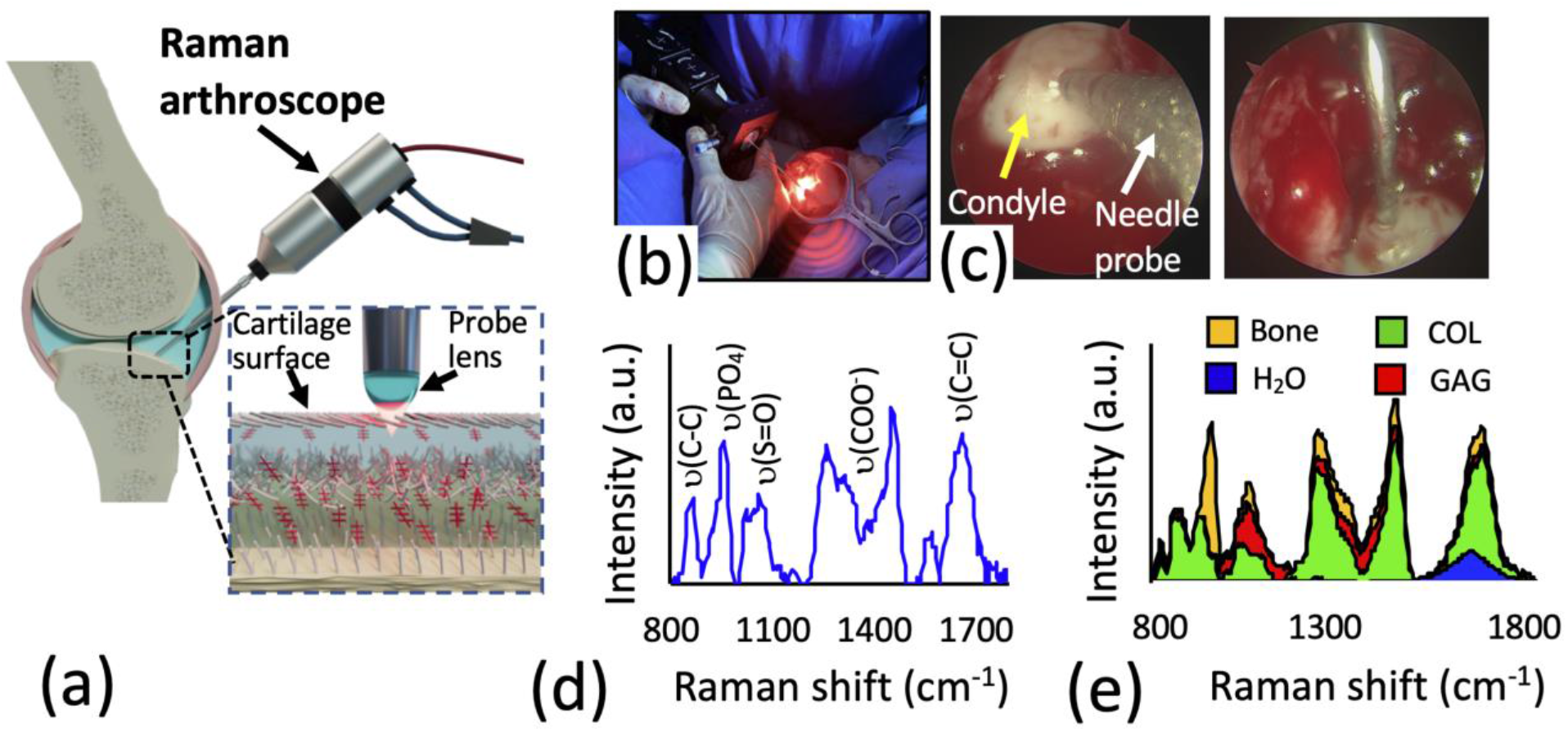
*In vivo* Raman arthroscopy diagnostics. **(a)** Schematic of fiber-optic Raman needle arthroscopy probe for OA diagnostics. The Raman spectroscopy system consists of a near infrared (NIR) laser, a spectrometer with an NIR deep depletion CCD, and a novel needle Raman probe that enables simultaneous acquisition of both the parallel and perpendicular polarized Raman signal. **(b)** *In vivo* Raman spectroscopy assessment of ovine stifle joint. **(c)** Raman probe in direct contact with femoral condyle as visualized by arthroscopic camera. **(d)** Raman spectra acquired *in vivo* of ovine femoral condyle articular cartilage. **(e)** 2D stacked area graph showing contribution of GAG, COL, H_2_O, and subchondral bone to composite ovine cartilage Raman spectra after multivariate linear regression (described in *Methods*).

### Multivariate Statistical Analysis

The Raman spectral contribution of GAG and water are characteristically “buried” under the much stronger COL signal (Fig.S1), thereby obscuring assessment of tissue GAG and H_2_O content. Here, we build upon our prior work on multivariate spectral analysis by decomposing and isolating the relative contribution of the major cartilage ECM constituents (GAG, COL, and H_2_O) to the Raman cartilage spectra using regression coefficients derived from the multivariate least-squares-regression analysis:

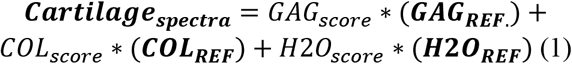

Where ***GAG_REF_**, **COL_REF_***, and ***H_2_O_REF_*** are the component spectra of purified reference chemicals of each ECM constituent (Fig.S1). The *GAG_score_, COL_score_*, and *H_2_O_score_* “scores” are the regression coefficients that reflect the contribution of the spectra of each constituent element to the cumulative Raman cartilage spectra.

### Raman probe GAG measurements *ex vivo*

The capability of our Raman arthroscopic probe and multivariate analysis to portray cartilage GAG content was assessed. Capitalizing on low ECM compositional heterogeneity ^24^, deep zone cartilage was extracted from explants (∅5mm±0.8mm) of femoral condyle hyaline cartilage of 2-month-old calves (Green Village Packing Co, NJ; N=5 animals). To simulate the progressive loss of GAG, as observed in OA, explants were subjected to step-wise GAG depletion using timed exposure (0, 4, 24, or 48 hours; n=10 explants per group) of 4M guanidine hydrochloride (GuHCl). Overnight exposure to 3mg/mL hyaluronidase (HA-dase; 37°C and pH 6.0) produced full GAG depletion. Raman spectra were acquired on the central region of each explant using a convex lens and compared to the GAG, COL, and H_2_O content and equilibrium compressive Young’s modulus (E_Y_) of a ∅3mm central core.

### Raman probe depth selective measurements *ex vivo*

As cartilage degeneration occurs in a depth-dependent manner, initiating predominantly in the topmost regions of the tissue, we next investigated how depth selectivity affects the quantification of GAG using a shallow focusing lens (needle and ∅2 mm ball lens) and deep focusing lens (convex lens) (Fig.S2). To induce progressive depth dependent GAG reduction, mimicking OA progression, full thickness bovine cartilage explants (∅6mm) were treated with 500μg/mL trypsin (pH 7.2 at 4°C) for 0, 0.5, 2, 4, or 8 hours (n=4 explants per group). Raman spectra were acquired at the articular surface using the ball lens and convex lens. Subsequently, explants were fixed, paraffin-embedded, sectioned, and stained with Safranin-O/Fast Green. Safranin-O colorimetric profiles were mapped through the depth using the red channel intensity and normalized to the average intensity at the 1.5mm depth position. For each lens, profiles were multiplied by the DOP decay curves (see *Results*) and integrated through the tissue depth, yielding the colorimetric-based GAG content within the lens-specific Raman measurement window. For each lens, colorimetric GAG was compared to Raman GAG scores. On a separate batch of trypsin-digested explants, the indentation elastic modulus of the cartilage surface was compared to Raman GAG scores acquired with the ball lens.

### Raman probe GAG measurements in human cartilage *ex vivo*

To establish the clinical relevance of the derived Raman GAG scores, Raman needle probe measurements were performed on ∅4.0mm chondral explants (n=13) excised from the distal femoral condyles of three cadaveric human knees (NDRI; age/sex: 70/♀, 75/♂, 65/♀; 4-5 explants per donor). Specimens exhibited an average thickness of 1.1±0.4mm and no visual signs of surface damage or fibrillation (Outerbridge scores 0-1). Raman spectra were acquired with both the ball lens and convex lens. Subsequently, explants were diametrically cut in half. For one half, the topmost 500μm of cartilage was excised for GAG content analysis. Histological sections from the other half were analyzed for a modified-Mankin-based Safranin-O-Fast-Green (SOFG) stain uptake score, based on percentage depletion per total area of unmineralized articular cartilage, as described ^25^. Ball lens Raman GAG scores were compared to surface cartilage GAG content measures. Convex lens Raman GAG scores were compared to the SOFG stain uptake score. To illustrate Raman arthroscopy’s ability to characterize the spatial variation in tissue composition along a contiguous joint surface, Raman GAG scores were acquired at discrete anatomical sites along the articular surface of an excised human femoral head specimen, obtained from a total hip arthroplasty procedure (50/♀; Kellgren-Lawrence grade 1). Raman GAG scores were acquired at three discrete anatomic sites along the articular surface and compared to corresponding Safranin-O intensities.

### Polarized Raman probe assessment of zonal collagen alignment *ex vivo*

We evaluated whether our polarized Raman probe could evaluate loss of the SZ during cartilage degeneration. Full thickness bovine explants were subjected to mechanical surface abrasion to remove sequential zonal layers (n=5 explants per group): no abrasion (SZ intact), mild abrasion (exposing the MZ), and severe abrasion (exposing the DZ) (detailed methodology in Supplementary Material). The parallel and perpendicular polarized Raman spectra (n=5) were collected from each sample. The depolarization ratio (perpendicular / (parallel + x)) of each spectral set was calculated and used for input to a partial least squares–discriminant analysis (PLS-DA), with leave-one-out cross validation. Orthogonal latent variables were derived from Raman intensity peak positions highly associated with polarization sensitive collagen bands that maximized the covariance between spectral variation and abrasion group affinity.

### Raman probe cartilage thickness measurements *ex vivo*

For Raman thickness assessments, the cartilage layer of bovine osteochondral explants was variably excised to achieve chondral thicknesses ranging from 0.3mm to 2.1mm. Raman spectra were acquired via the convex lens to better detect the Raman spectra from the subchondral bone through diffuse light scattering. Using known reference spectra of cartilage ECM (GAG, COL, H_2_O) as well as bovine subchondral bone (***Bone_REF_***; Fig.S3) and its regression coefficient (*Bone_score_*), multivariate linear regression was applied to the aggregate Raman spectra.

### Inter-anatomical variability as revealed by Raman spectroscopy *ex vivo*

We further assessed the capability of Raman biomarkers to predict properties of cartilage from different anatomical sites of the ovine knee joint, which exhibit biochemical and mechanical variability ^26^. Raman probe measurements and mechanical testing were performed on osteochondral explants (∅6.0mm) from the medial and lateral femoral condyles, and tibial plateaus (1-2 explants per surface) of a skeletally mature ovine knee joint. The chondral layer of explants was subsequently analyzed for GAG content, H_2_O content, and thickness. Univariate and multivariate regression using the compositional Raman scores was applied to predict biochemical contents, thickness, and elastic modulus.

### Raman arthroscopy for *in situ and in vivo* diagnostics

Confounding factors related to *in situ* intra-articular Raman diagnostic measurements were further assessed, including: 1) interference from synovial fluid (SF), 2) sensitivity of Raman biomarkers to probe-to-cartilage surface incidence angle, and 3) acquisition time to achieve reliable Raman signal (all described in *Supplementary Material*). Further, *in situ* measurements of cartilage ECM composition were performed on intact *ex vivo* bovine antebrachiocarpal (wrist) joints before and after intra-articular enzymatic GAG depletion treatment using the Raman arthroscopy probe inserted intra-articularly through a 10-gauge hypodermic needle trocar (*Supplementary Material*).

*In vivo* Raman arthroscopy was approved by University of Pennsylvania Veterinary School IACUC. Raman spectra were collected from the distal femoral condyle articular cartilage of a live skeletally-mature sheep via a mini arthrotomy of the stifle joint.

## Results

### Raman probe GAG measurements *ex vivo*

Chemical treatments induced a stepwise depletion of GAG from cartilage explants (Fig.2a). COL content (mean 4.4±0.9% per wet weight [%ww]) and H_2_O content (mean 86.7±2.6%ww) were minimally altered by these treatments. Concomitant with chemically-induced GAG depletion was a prominent decrease in Raman signal intensity in the ranges 1000-1100 cm^-1^ and 1200-1300 cm^-1^ (Fig.2b). Following multivariate regression analysis (equation 1), the Raman GAG score decreased in proportion to the reduction of GAG from the explants; the Raman scores for COL and H_2_O were less affected (Fig.2c). The cumulative spectral contribution of the individual ECM constituents accounted for 94% of the variation of the composite cartilage spectra (Fig.2d; R^2^=0.94±0.01; p<0.001). The GAG scores predicted 95% of the variation in measured tissue GAG content (Fig.2e; R^2^=0.95; p<0.001; Table S1) and 74% of the variation in the measured compressive modulus (E_Y_) for all explants (Fig.2f; R^2^=0.74; p<0.001), demonstrating the capacity of our Raman probe to predict progressive GAG loss and mechanical softening of hyaline cartilage associated with OA.

**Fig 2.**
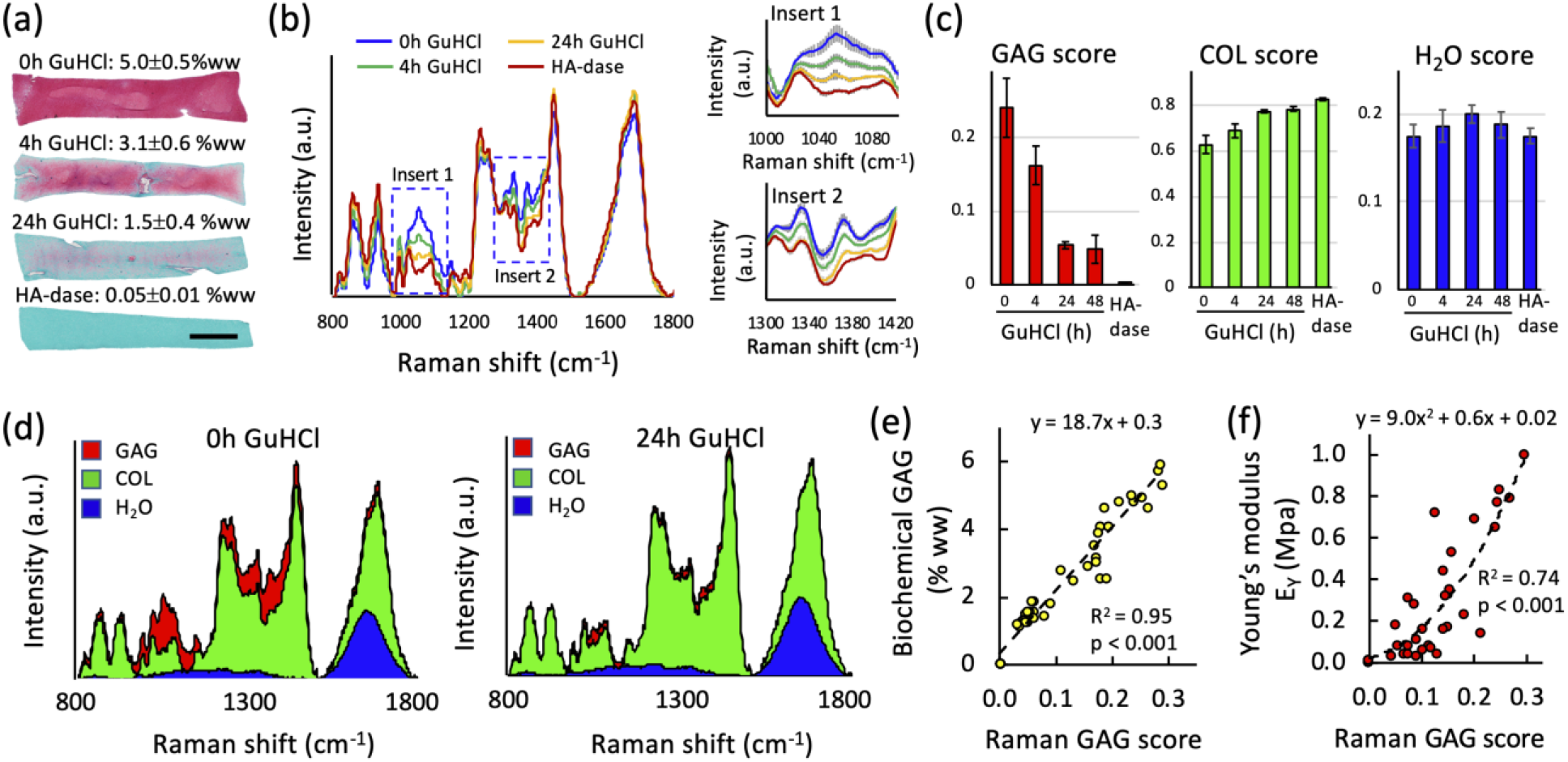
Raman probe GAG measurements. (**a**) Guanidine hydrochloride (GuHCl) and hyaluronidase (HA-dase) induced GAG depletion of cartilage explants (Safranin-O histology and DMMB-measured GAG levels). Scale bar = 1mm. (**b**) GuHCl/HA-dase-induced decrease of measured Raman probe spectra intensity at 1000-1100 cm^-1^ and 1300-1450 cm^-1^ wavenumbers (mean ± standard deviation). (**c**) Regression coefficients (scores) for GAG, COL, H_2_O from multivariate linear regression decomposition of Raman spectra for GuHCl/HA-dase timed exposure groups. (**d**) 2D stacked area graph showing cumulative contribution of GAG, COL, H_2_O spectra to composite Raman cartilage spectra after multivariate linear regression decomposition. GAG spectral contribution attenuated after GAG depletion by GuHCL, while COL and H_2_O contributions relatively unaffected. Bivariate regression between Raman GAG scores vs: (**e**) assay-measured GAG content and (**f**) compressive Young’s modulus (E_Y_) for explants.

### Raman probe depth selective measurements *ex vivo*

The ball lens and convex lens exhibited different DOP values as verified by the Raman signal attenuation of polystyrene substrates under varying thickness cartilage layers (Fig.3a). Cartilage explants exhibited progressive surface GAG depletion with increasing trypsin treatment time (Fig.3b). Raman GAG scores deduced from Raman spectra obtained through the surface-targeting ball lens predicted 86% of the GAG tissue content (Fig.3c; R^2^=0.86; p<0.001; Table S2). However, Raman GAG scores deduced from spectra obtained through the deep focus convex lens, only predicted 40% of the GAG content (R^2^=0.4; p<0.001; Table S2); GAG scores were influenced by residual GAG remaining in the deep zone, due to restricted diffusion of the GAG-depleting enzyme. Raman GAG scores from the ball lens further predicted 86% of the elastic modulus (Fig.3d; R^2^=0.86; p<0.001). These results indicate that Raman arthroscopy with a surface-focusing lens is advantageous for quantification of cartilage surface GAG depletion and softening.

**Fig 3.**
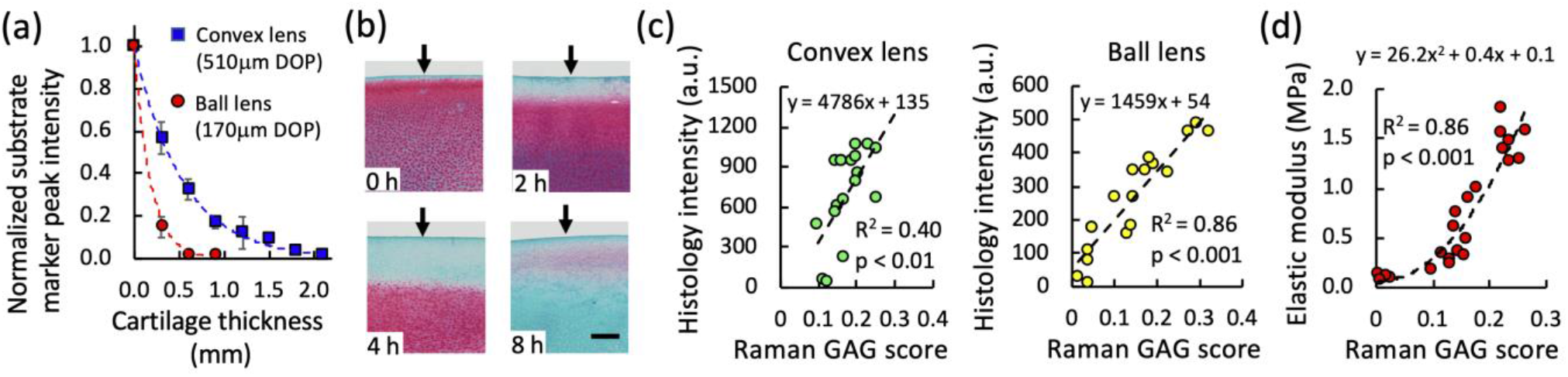
Raman probe depth selective measurements. (**a**) Lens-specific depth of penetration (DOP) based on measured decay of polystyrene substrate Raman signal under variable cartilage thickness layers (Supplementary Data). (**b**) Representative Safranin-O sections depicting trypsin-induced GAG depletion from the articular surface of cartilage explants. Scale bar = 250μm. (**c**) Bivariate linear regression between Raman probe-measured GAG scores and colorimetric Safranin-O-measured GAG content for deep focusing convex lens and surface-targeting ball lens. (**d**) Bivariate linear regression between ball lens measured GAG scores and indentation elastic modulus.

### Raman probe GAG measurements in human cartilage *ex vivo*

Human chondral explants exhibited a range of Safranin-O staining intensities and GAG contents (Fig.4a). The ball-lens-acquired GAG scores predicted 66% of the variation of GAG content in the topmost 500□m tissue layer (Fig.4b; R^2^=0.66; p<0.001; Table S3). Convex-lens-acquired GAG scores predicted 53% of the variation of a Safranin-O-Fast-Green (SOFG) stain uptake score (Fig.4c; R^2^=0.53; p<0.01; Table S3). Raman GAG scores were acquired at discrete anatomic sites along the articular surface of an excised human femoral head (Fig.4d-e). Low Raman GAG scores were observed in the GAG-depleted hyaline cartilage, comprising the loaded region of the hip joint, while high GAG scores were observed in the GAG-replete hyaline cartilage comprising the unloaded region of the hip joint.

**Fig 4.**
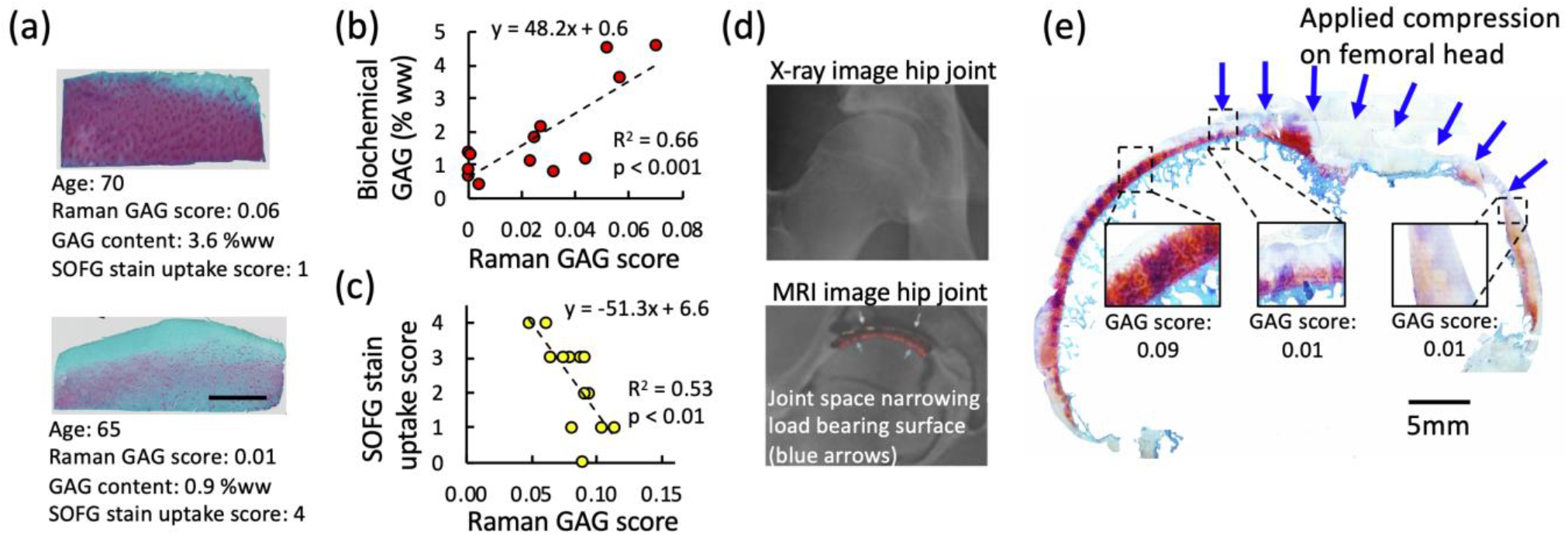
Raman probe GAG measurements in human cartilage e*x vivo*. (**a**) Representative Safranin-O histological sections of GAG-replete and GAG-depleted cartilage explants from human autopsy donors along with Raman GAG scores, GAG content, and SOFG stain uptake score. Scale bar = 1mm. (**b**) Bivariate linear regression between Raman probe GAG scores (ball lens measured) and DMMB-measured GAG content of n=13 human explants *ex vivo*. (**c**) Bivariate linear regression between Raman probe GAG scores (convex lens measured) and SOFG stain uptake scores. (**d**) Representative radiographic and MRI sagittal plane images of arthritic hip joint, illustrating joint space narrowing at superior femoral head, corresponding to load bearing region of hip during standing. (**e**) Raman probe GAG scores acquired at discrete anatomic regions along a sagittal slice of a human femoral head articular joint surface *ex vivo*. GAG scores reflect depletion of GAG and cartilage thinning observed on MRI and histological section.

### Polarized Raman probe for assessment of zonal collagen alignment

Consistent differences between the perpendicular and parallel polarization spectra were observed among the groups (no abrasion, mild abrasion, and severe abrasion) at Raman intensity peak positions highly associated with polarization sensitive collagen bands: 861, 930, 1257, 1448 and 1654 cm^-1^ (Fig.5a) ^27^. The depolarization ratio spectra (i.e., intensity ratio between the perpendicular and parallel components of the Raman scattered light) revealed that differences in the polarized Raman spectra portrayed the varied zonal organization of the collagen fiber network (Fig.5b). From PLS-DA, latent variable (LV) loadings LV1 and LV2 demonstrated good discriminatory separation among the abrasion groups (Fig.5c) and incorporated diagnostically relevant spectral variations that reflected the integrity of the SZ collagen network (LV1: 10.36% and LV2: 6.48%; Fig.5d). Compared to LV1, LV2 better differentiated samples where the SZ was intact (p < 0.001), therefore LV2 was used as a Raman collagen alignment factor (RCAF) to depict the extent that SZ collagen was retained (Fig.5e). These results show that collagen alignment and SZ degeneration can be quantified by taking advantage of the polarization response of cartilage.

**Fig 5.**
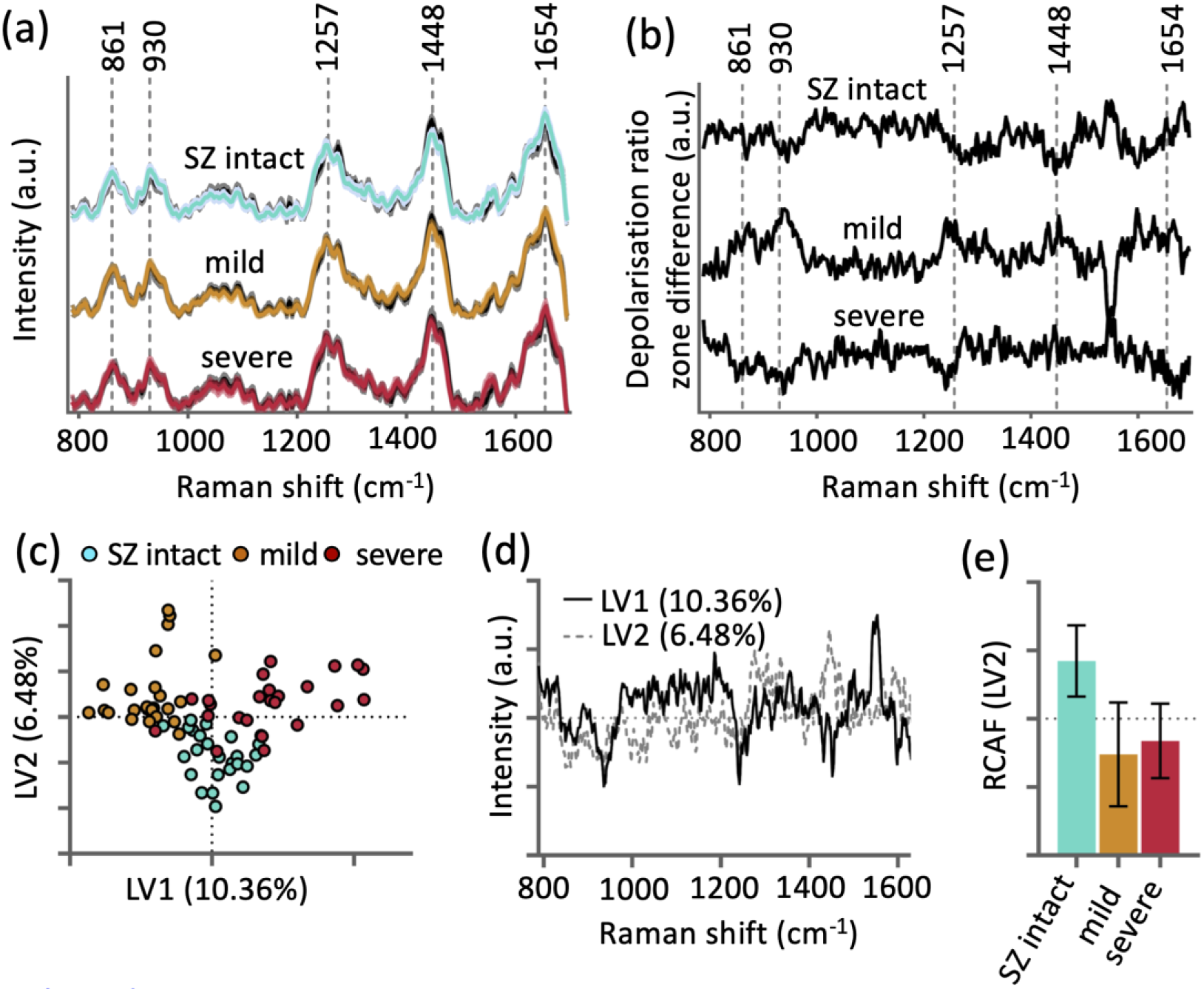
Polarized Raman probe_collagen alignment measurements. (**a**) Mean polarized Raman spectra (perpendicular [color lines] and parallel [black lines]) of cartilage with an intact SZ, mild abrasion and severe abrasion of the surface layer. (**b**) Difference spectra of the polarization ratio (parallel/perpendicular) reveal differences in Raman spectra due to integrity of SZ collagen. (**c**) Partial least squares discriminant analysis (PLS-DA) scores showing good separation of the different erosion groups. (**d**) PLS-DA latent variable (LV) loadings LV1 and LV2. (**e**) Raman collagen alignment factor (RCAF) for detecting extent of SZ abrasion corresponding to LV2.

### Raman probe cartilage thickness quantification

Following multivariate regression analysis of the Raman spectra of variable thickness bovine osteochondral explants, the contribution of bone signal to the aggregate spectra decreased with increasing cartilage thickness (Fig.6a). The regression coefficient for the subchondral bone contribution (bone score) to the cumulative Raman spectra varied inversely with thickness of the chondral layer, following an exponential decay function (Fig.6b). The bone score predicted 90% of the variability in cartilage thickness (Table S4).

**Fig 6.**
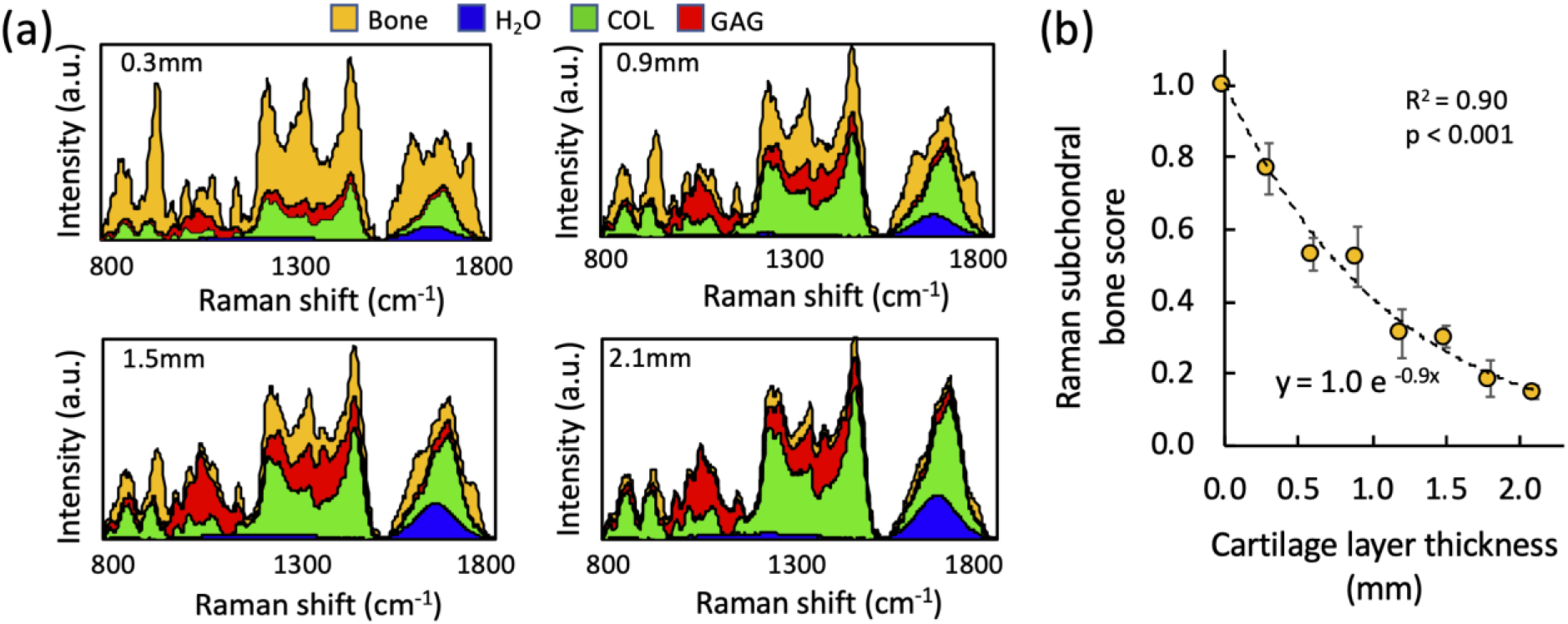
Raman probe cartilage thickness measurements. (**a**) 2D stacked area graph showing contribution of GAG, COL, H_2_O, and subchondral bone to composite Raman cartilage spectra after multivariate linear regression decomposition for osteochondral cartilage explants with a 0.3, 0.9, 1.5, and 2.1 mm thick cartilage layer. (**b**) Bivariate regression between Raman probe measured subchondral bone score versus cartilage layer thickness.

### Inter-anatomical variability as revealed by Raman spectroscopy *ex vivo*

For ovine osteochondral explants (Fig.7a), the Raman GAG score predicted 96% of the GAG content (p<0.001; Fig 7b), H_2_O score predicted 85% of the water content (p=0.01; Fig.7c), and bone score predicted 78% of the thickness (p=0.02; Fig.7d). While GAG score alone predicted 80% of indentation modulus (p=0.02; Fig.7e), multivariate regression using a linear combination of the Raman scores for GAG and H_2_O predicted 94% of the indentation modulus (p<0.01; Fig.7f)

**Fig 7.**
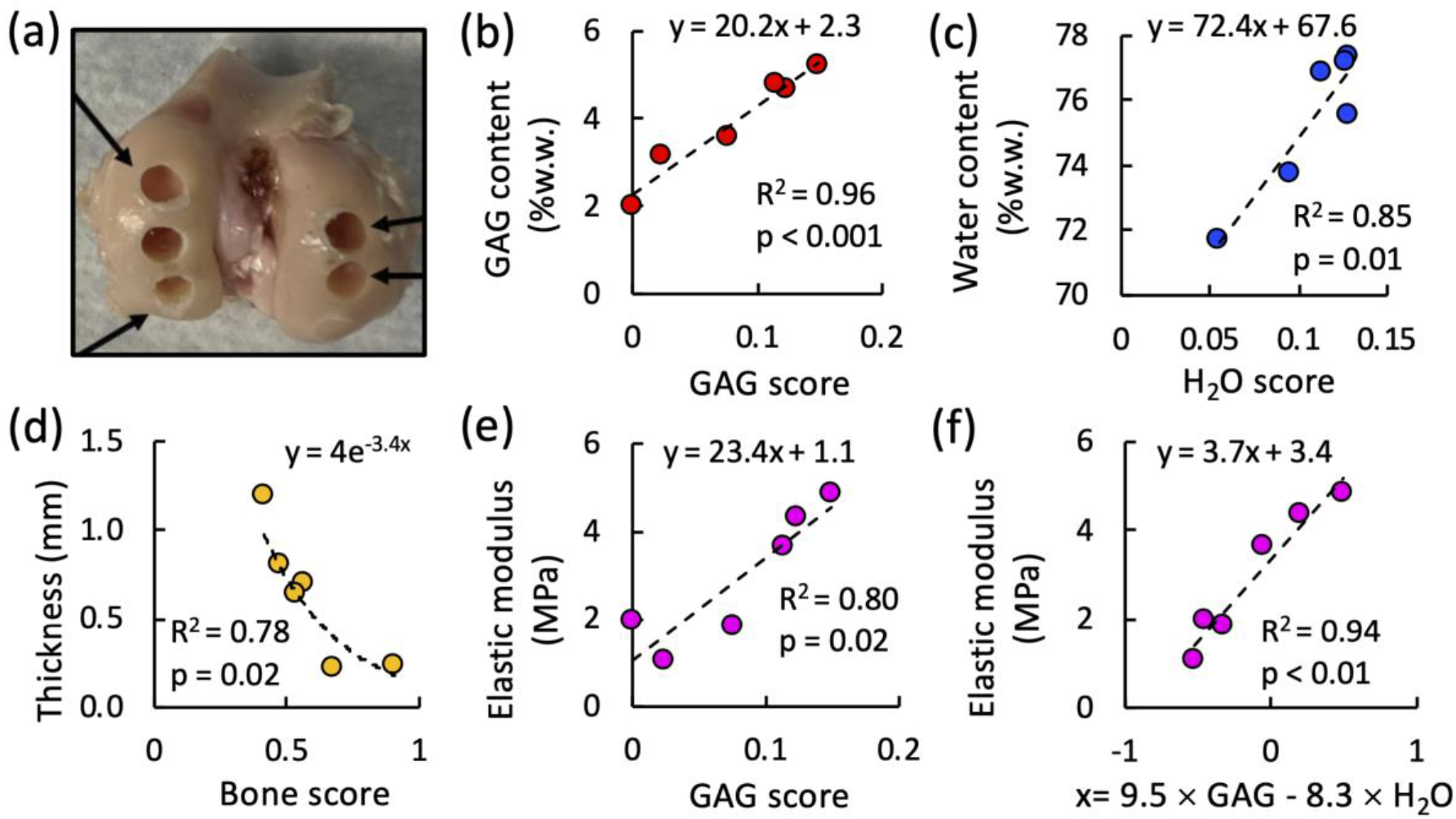
Raman probe inter-anatomical measurements of cartilage *ex vivo*. (**a**) Representative cored regions on ovine femoral condyle articular surfaces. Bivariate regression between (**b**) Raman GAG score and GAG content, (**c**) Raman H_2_O score and water content, (**d**) Raman bone score and cartilage thickness, (**e**) Raman GAG score and elastic modulus. (**f**) Multivariate regression between linear combination of Raman GAG and H_2_O scores (9.5×GAG_score_-8.3×H_2_O_score_), and elastic modulus.

### Raman arthroscopy *in situ* and *in vivo* diagnostics

Assessments of potential confounding factors related to *in situ* Raman arthroscopy measurements demonstrated that: 1) Raman biomarkers were insensitive to the presence of SF (Fig.S4a), 2) 20°variation from a normal (90°) probe incidence angle to the cartilage surface had no significant effect on Raman biomarkers (p<0.05; Fig.S4b), and 3) integration time did not significantly affect measured Raman biomarkers (Fig.S4c), indicating Raman arthroscopy measurements obtained in as little as 0.5 seconds will not compromise diagnostic capability. Further, for intra-articular Raman arthroscopy assessments on intact wrist joints, Raman GAG scores were reduced by 75% after trypsin treatment corresponding to 86% reduction in cartilage GAG content (Fig.S5).

The distal femoral condyle articular cartilage of a live skeletally-mature sheep was accessed via a mini arthrotomy of the stifle joint (Fig.1). The Raman needle probe was placed in gentle contact with the articular surface of the femoral condyle under image guidance. Raman spectra were acquired over a 10 second integration time to obtain the highest SNR for this *in vivo* demonstration. High quality Raman spectra were acquired *in vivo* (Fig.1d), akin to *ex vivo* measures. The cumulative spectral contribution of the individual ECM constituents and subchondral bone derived by multivariate regression analysis accounted for 86% of the variation in the composite spectra (Fig.1e; Raman scores: GAG=0.16, COL=0.58, H_2_O=0.11, subchondral bone=0.10; R^2^=0.86; p<0.001). *In vivo* scores were similar to those obtained *ex vivo* on ovine osteochondral explants (Fig.7). This demonstration highlights the feasibility of *in vivo* Raman arthroscopy by demonstrating successful compositional quantification *in vivo* and the ability to maneuver the needle probe in a surgical setting.

## Discussion

We present a novel needle-based arthroscopic platform for achieving real-time, polarized Raman spectroscopy quantification of changes in cartilage composition, structure, and material properties associated with OA. Using *ex vivo* bovine models of OA, aging human cartilage explants, and anatomical-site-varied ovine cartilage explants, we demonstrate that our Raman arthroscopic probe can measure biomarkers that accurately predict changes in GAG content and collagen disorganization associated with the degradation of hyaline cartilage in OA. A critical innovation, described herein, is the implementation of multivariate regression to ascertain the contribution of individual spectra corresponding to the ECM constituents GAG, COL, H_2_O and bone to the cumulative Raman cartilage spectra ^24,28^. The derived regression coefficient biomarkers (GAG scores) account for 95% of the variation of the GAG content of enzymatically depleted bovine explants, 66% of the GAG variation in human chondral explants, and 94% of the GAG variation in ovine osteochondral explants. Additionally, the derived Raman H_2_O and COL scores can reveal damage to the integrity of the tensile collagen matrix that give rise to cartilage swelling in OA ^29^. Raman biomarkers can further be used to predict cartilage mechanical properties. Raman GAG scores account for 86% of the variation of the elastic modulus of GAG depleted bovine explants. For healthy native ovine cartilage specimens, the multivariate combination of Raman-derived biomarkers (GAG and H_2_O score) accounts for 94% of the cartilage elastic modulus. This demonstration highlights the important capability of composite Raman biomarkers to provide assessments of tissue functional material properties.

A potential limitation of this analysis is that purified reference chemicals may exhibit spectral profiles that are different from ECM *in situ*, thus contributing to variance in regression models. Future models may improve Raman assessments by accounting for the compositional and organizational complexities of ECM constituents *in situ*. The lower GAG score correlation exhibited for human specimens likely results from the spatial disparity between the tissue regions of interest analyzed with the Raman probe and biochemical assays—an effect that is likely more pronounced for these heterogeneous specimens. Future validations may benefit from the use of direct ECM measurement techniques with higher spatial resolution that can offer more faithful comparisons with biomarkers acquired within the penetration depth of the Raman probe.

To further evaluate the zone-dependent, anisotropic microstructure of the collagen matrix, we incorporated polarized Raman spectra into our platform to exploit differences between the perpendicular and parallel polarization spectra. This novel functionality, achieved by tightly focusing the distal ball lens to avoid bulk tissue polarization scrambling, enables assessment of the alignment and organization of the SZ collagen matrix. To maximize diagnostically relevant spectral variations that reflect the integrity of the SZ collagen, PLS-DA was applied to the depolarization spectra, giving rise to latent variables (LV1, LV2) that incorporated Raman intensity peak positions highly associated with polarization sensitive collagen bands. LV2 efficiently discriminates an intact SZ and can therefore be used as an alignment factor to depict the extent that SZ collagen was retained.

While the diagnosis of surface GAG and collagen loss requires utilization of a ball lens that collects Raman signal predominantly from the topmost cartilage regions, Raman cartilage thickness measurements may require a deep focusing lens to collect sufficient Raman signal from the subchondral bone. Thus, the lens configuration selected depends on the diagnostic metric to be quantified. For immature bovine specimens, to resolve Raman-based cartilage thickness, we used a large, non-needle-based lens to target the subchondral bone. In future iterations of our device, an interchangeable deep tissue lens (hemispherical lens or custom manufactured high DOP micro lens) will be incorporated into the needle probe.

To demonstrate the feasibility of translating Raman needle arthroscopy to the clinic, we acquired high quality Raman spectra of articular cartilage comprising the ovine femoral condyle *in vivo*, which proved to be similar to *ex vivo* assessments. Clinical translation is further supported by establishing: 1) Raman spectra can be obtained in as little as 0.5 seconds, 2) the derived Raman GAG scores are insensitive to the presence of SF, and 3) the probe incidence angle can vary up to 20°from normal to the cartilage surface without compromising diagnostic accuracy. Further, *in situ* intra-articular Raman spectra acquired on intact bovine diarthrodial joints before and after enzymatic-induced degradation validated that Raman GAG scores were consistent with measured GAG content.

There has been growing interest in Raman spectroscopy as a potential OA diagnostic technology, building upon prior work on IR spectroscopy ^30-32^. Prior *ex vivo* studies have examined Raman spectral changes in explanted late stage OA cartilage tissues ^20-22,33^ or cartilage subjected to mechanical damage ^16-18^. With the exception of Unal et al. ^33^, who used high-wavenumber Raman peak ratios to measure cartilage water content but not GAG or collagen, prior Raman assessments have consisted of univariate peak analysis or principle component analysis (PCA), which do not provide quantification of biochemical or structural tissue changes relevant to the tissue’s functional performance. Further, with the exception of Esmonde-White et al. ^19^, which used a probe to measure cartilage erosion, previous Raman investigations have been performed on benchtop microscopy systems that are incompatible with *in situ* intra-articular Raman evaluations. Our study represents the first *in vivo* Raman diagnostic investigation to utilize a clinically compatible needle arthroscopic probe to measure biomarkers that are associated with the pathognomonic changes of OA: GAG depletion, SZ collagen loss, erosion (thinning), and swelling (increased hydration). In future work, we can incorporate additional key molecular constituents into spectral decomposition models (e.g., GAG/COL subtypes, lipids, crosslinks, DNA), thus providing additional biomarkers to potentially improve diagnostics on specimens with increased compositional complexity.

This work supports the use of our Raman probe as a transformative diagnostic platform for objective monitoring of treatment outcomes of emerging OA therapies. It can be employed over the hierarchy of cartilage tissue model systems, including: 1) *in vitro* non-destructive, repeated-measure assessments of the efficacy of novel OA therapeutics in ameliorating degeneration of live explant tissues, and 2) *in vivo* assessments (preclinical and clinical) as a minimally invasive, real-time diagnostic tool for articular cartilage disease stage and evaluation of treatment response. Future *in vivo* investigations will be required to directly ascertain the capability of Raman arthroscopy to monitor degeneration and repair. Raman arthroscopy can complement other state-of-the-art and emerging OA diagnostic platforms (MRI (T1ρ, dEGEMRIC) ^34,35^, ultrasound ^36^, contrast enhanced CT ^37^, OCT ^38^) in portraying changes in cartilage composition, structure, and material properties. While the current diagnostic gold-standard, MRI, is non-invasive and can image the entire joint, it is limited by spatial resolution of the cartilage surface ^39^, systemic administration of potentially toxic contrast agents ^40^, extended imaging times, expense, lack of portability, and/or infrastructure requirements. Alternatively, Raman arthroscopy can serve as a low cost, minimally invasive, portable diagnostic platform that achieves rapid and safe (no radiation or toxic contrast agent exposure) assessments of cartilage composition. While Raman arthroscopy is limited to point-based measures, it can be interfaced with image guidance (e.g., Arthrex NanoScope) to achieve targeted assessments of specific anatomical sites. Raman biomarkers appear to be more sensitive to changes in cartilage composition than MRI (T1ρ relaxation times account for approximately 20% of cartilage GAG variation ^41^), although direct comparisons will need to be performed in future work. As such, Raman can potentially lead to the development of clinical trials that incorporate fewer participants, shorter durations, and patient cohorts with earlier stages of OA (before irreversible degeneration has transpired)—together, improving the likelihood of the identification of chondroprotective OA therapies. Raman arthroscopy may further provide a unique utility for diagnostics of joints with characteristic thin cartilage layers (e.g., shoulder, elbow) where image-based cartilage evaluation is particularly limited.

In the future, Raman needle arthroscopy can be performed as a cost-effective, point-of-care office procedure capable of diagnosing the early stages of OA before irreparable changes in cartilage biochemical and biophysical properties are evident radiographically. Raman arthroscopic screening can be performed on patient populations known to be at high risk for developing OA as a consequence of traumatic joint injury, physical occupations, internal derangement, obesity, joint malalignment, and genetic predisposition. The periodic monitoring of cartilage biomarkers may allow for the timely prescription of emerging chondroprotective therapies, such as biologics, viscosupplements, lifestyle changes (weight loss, activity cessation), physical therapy, or surgical reconstruction. As such, our Raman arthroscopy platform can set the foundation for the identification of novel OA therapies and subsequently support their administration in the clinic.

## Acknowledgments

**General**: We thank Dr. Thomas Schaer for assistance with in vivo ovine measures, Joshua Auger for assistance with procurement and processing of human cartilage specimens, and Sedat Dogru and Chenhao Yu for their assistance in implementing mechanical testing protocols.

## Author contributions

M.B.A., M.S.B., B.D.S, M.W.G designed and supervised the study, interpreted the data, and wrote the paper. K.K., M.W., J.Z., W.W. performed Raman measurements, analyzed Raman data, performed biochemical analysis. M.J. performed polarized Raman measurements. J.B.E. performed histological analyses. All authors contributed to scientific discussions and preparation of the manuscript.

