## Supplementary Data for "Raman Needle Arthroscopy for *In Vivo* Molecular Assessment of Cartilage"

### Raman needle arthroscopy Instrumentation

A custom polarized Raman needle arthroscopy system was developed for *in vivo* OA diagnostics by intra-articular entry through a hypodermic needle (Fig.1a-c). We utilize a hollow needle and avoid the use of distal fiber-optics in order to fully eliminate silica background signal, a critical aspect for clinical Raman database standardization and translation. The Raman needle arthroscopy system consists of a near infrared (NIR) laser, a spectrometer with an NIR deep depletion CCD, and a novel needle Raman probe that enables simultaneous acquisition of both the parallel and perpendicular polarized Raman signal (Jensen et al. Opt Lett. 2020; 45(10):2890-3). The laser (B&W CleanLaze 785 nm, 500 mW) is coupled to the Raman probe, where a polarization filter and a laser bandpass filter polarize and clean the laser light before it is directed through the needle probe (∅ = 2 mm, l = 50 mm) and tightly focused onto the tissue using a 2.0 mm sapphire ball lens. The Raman scattered light reflected from the superficial tissue is passed back through the needle, via a dichroic mirror, a long pass filter, and into a polarization beam splitter, where the Raman signal is split into two separate polarized components that are focused onto a bifurcated optical fiber. The two fibers are coupled to the spectrometer (Princeton Instruments LS785 and Pixis NIR deep depletion camera 400x1340 pixels), and the separation of the fiber cores creates two distinctly separated Raman spectra (perpendicular and parallel) on the CCD. The two CCD segments are hardware binned and read out simultaneously. Summation of the parallel and perpendicular polarization provides conventional Raman biochemical analysis while the depolarization ratio offers insight into the alignment and structure of the tissue.

### Raman arthroscopy spectral acquisition

The depth of penetration (DOP) of the Raman arthroscope was varied by incorporating a controlled focal length lens at the probe tip. Measurements were performed using either a ∅2.0mm ball lens (170µm DOP) integrated into a needle bore with the probe tip placed in gentile contact with and normal to the cartilage surface, or a larger convex lens (510µm DOP) with the focal point being on the cartilage surface. Raman spectra were acquired with a 10 second acquisition time to obtain highest possible SNR, unless otherwise noted. Spectra were processed by subtracting a premeasured background signal, then intensity calibrated using the standard NIST 2241 Raman spectra reference material. Subsequently, Raman spectra were wavelength calibrated using an atomic lamp (Ocean Optics HG-1). Background autofluorescence was removed using a constrained fifth-order polynomial fit function (Bergholt et al. Gastroenterology. 2014; 146(1):27-32). The Raman spectra were smoothed using a third order Savitzky-Golay filter (5 pixel window) and area-under-curve normalization.

**Raman spectra of cartilage ECM constituents**

To understand the spectral overlap of biomolecules in articular cartilage, Raman spectra were acquired of bovine cartilage explants and of the major ECM constituents (GAG, collagen [COL], water [$H_2O$]) in the tissue. ECM constituent spectra were generated from powdered purified reference chemicals of GAG (chondroitin-6-sulfate [Shark cartilage; Sigma C4384]) and COL (type-II collagen [chicken sternal cartilage; Sigma C9301]), and for $H_2O$ using phosphate buffered saline. Fig.S1 shows the significant overlap of the spectra of ECM constituents in cartilage, thus demonstrating the challenge of extracting quantitative biochemical information via univariate spectral analysis and supporting the need multivariate analytical platforms, as performed in this study. For multivariate regression analysis, lens-specific (ball lens or convex lens) $\boldsymbol{GAG_{REF}}$, $\boldsymbol{COL_{REF}}$, and $\boldsymbol{H2O_{REF}}$ reference component spectra were utilized.

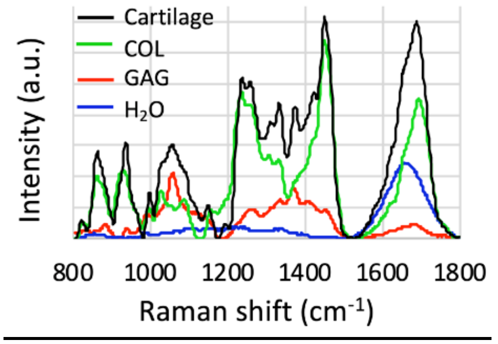


**Fig. S1.** Raman spectra of articular cartilage and its ECM constituents.

**Polarized Raman probe assessment of zonal collagen alignment *ex vivo*.** We evaluated whether our polarized Raman probe could evaluate loss of cartilage SZ during cartilage degeneration. Full thickness bovine explants were subjected to mechanical surface abrasion using a linearly reciprocating sander (600 grit) under 40kPa of normal stress. Abrasion was used to remove sequential zonal layers (n=5 explants per group): no abrasion (SZ intact), mild abrasion (120±26μm tissue removed, exposing the MZ), and severe abrasion (437±60μm tissue removed, exposing the DZ). All explants were GAG depleted via HA-dase to mimic OA GAG depletion. Using monochromic laser light excitation at a laser power of 140mW, the parallel and perpendicular polarized Raman spectra (n=5), were collected from each sample over a 5 second acquisition time. A total of 25 individual sets of polarized Raman spectra were measured for each group, resulting in 75 sets of polarized Raman spectra. The depolarization ratio (perpendicular / (parallel + x)) of each spectral set was calculated using the arbitrary DC component to avoid near infinite values. The depolarization ratio was used for input to a partial least squares–discriminant analysis (PLS-DA), with leave-one-out cross validation to build an unbiased model that discriminated among the surface abrasion groups. PLS-DA is a powerful multivariate regression technique that can efficiently extract the spectral variations of interest such as changes associated with alignment or chemistry. Orthogonal latent variables were derived from Raman intensity peak positions highly associated with polarization sensitive collagen bands that maximized the covariance between spectral variation and abrasion group affinity. All statistical analysis was performed with Matlab using PLS-Toolbox.

**Cartilage explant composition and material property measures**

The Young's modulus ($E_Y$) of GuHCl/HA-dase treated immature bovine cartilage explants (Fig.2f) was determined by measuring the equilibrium stress-strain response under 10% strain in unconfined compression. The indentation elastic modulus of the articular surface of trypsin-treated immature bovine cartilage explants (Fig.3d) and ovine osteochondral explants (Fig.7e-f) was determined via the Oliver-Pharr method (Oliver WC & Pharr GM. J Materials Research; 1992. 7: 1564-83) using the slope of the unloading curve after indentation with a 3mm hemispherical indenter under 100 □m displacement (loading/unloading ramp velocity 2µm/s). Explant water content was determined via gravitational measures before and after lyophilization. Explant GAG and COL contents were determined via dimethylmethylene blue assay and orthohydroxyproline assay, respectively, after proteinase-K digestion.

### Raman arthroscope lens depth of penetration

The depth a penetration of the Raman arthroscopic probe was modulated by incorporating a lens at the probe tip with a controlled focal length. Measurements were performed using either a ball lens (∅2.0mm sapphire, AWI Industries) or a larger convex lens (plano-convex N-BK7 ∅9.0 mm, f = 12.0 mm, AR Coating: 650-1050 nm, Thorlabs). The depth of penetration (DOP) of each lens, was approximated by measuring the Raman peak intensity of a polystyrene substrate under a varying thickness of bovine cartilage (Fig.S2a-b). The DOP was measured from the cartilage layer thickness at which the normalized polystyrene substrate 988 $cm^{-1}$ peak intensity decayed to a value of 0.37, (Fig.S2c), yielding a DOP value of 170□m for the ball lens and 510□m for the convex lens.

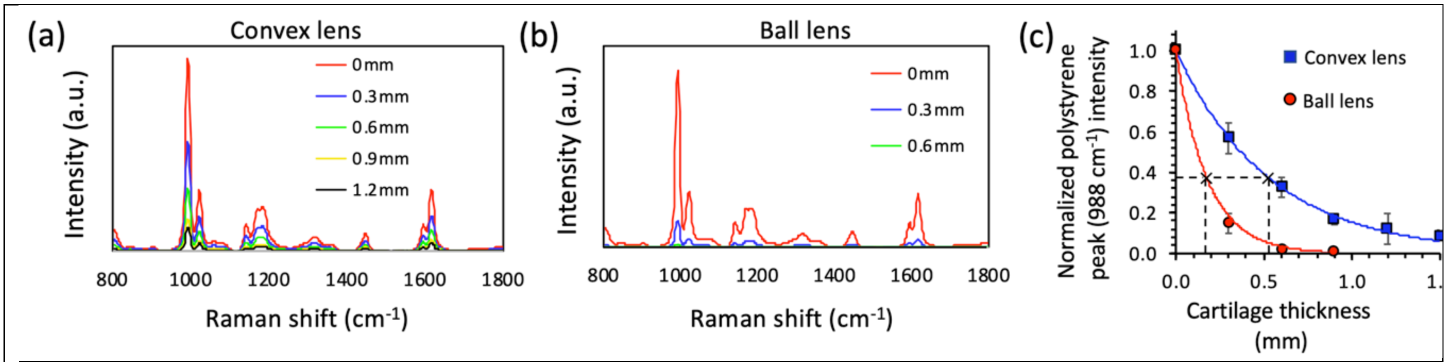


**Fig.S2**: Raman spectra of a polystyrene substrate under a varying thickness of cartilage as measured by (**a**) convex lens and (**b**) ball lens. (**c**) Intensity of polystyrene 988 $cm^{-1}$ under varying cartilage thickness (normalized to 0mm level) for convex and ball lens. "×" marks lens-specific cartilage thickness at which normalized peak intensity decays to a value of 0.37, as used to calculate lens DOP.

**Raman subchondral bone**

For Raman cartilage thickness measurements, multivariate linear regression analysis was performed with the addition of a subchondral bone term, incorporating the component reference spectrum of subchondral bone from biological specimens. Bone reference spectra were acquired using our probe from intact subchondral bone specimens cartilage layer completely excised from immature bovine femoral condyles (Fig.S3a) and skeletally-mature ovine femoral condyles (Fig.S3b). The bone reference spectrum for human cartilage was acquired from the human femoral head specimen (Fig.4e). The subchondral bone spectrum represents a combination of molecular species, predominantly hydroxyapatite (Fig.S3d) and lipids (the triglyceride triolein [Fig.S3e]) and with a minor contribution of collagen. Distinct peaks originate from hydroxyapatite at 950cm$^{-1}$ and lipids (ester vibrations at 1745 cm$^{-1}$, ν(C=C) at 1650cm$^{-1}$, $CH_2$ deformations at 1435cm$^{-1}$). The collagen contribution to the spectra is largely absent, or buried, as evidenced by a lack of common protein peaks, such as phenylalanine at 1004 cm$^{-1}$. The immature bovine subchondral bone specimen shows an additional strong contribution of blood in the tissue, which is characteristic of immature subchondral bone, as evidenced by heme porphyrin ring vibrational bands (near 1340 and 1600 cm$^{-1}$) (Lemler et al. Anal Bioanal Chem; 2014; 406(1)).

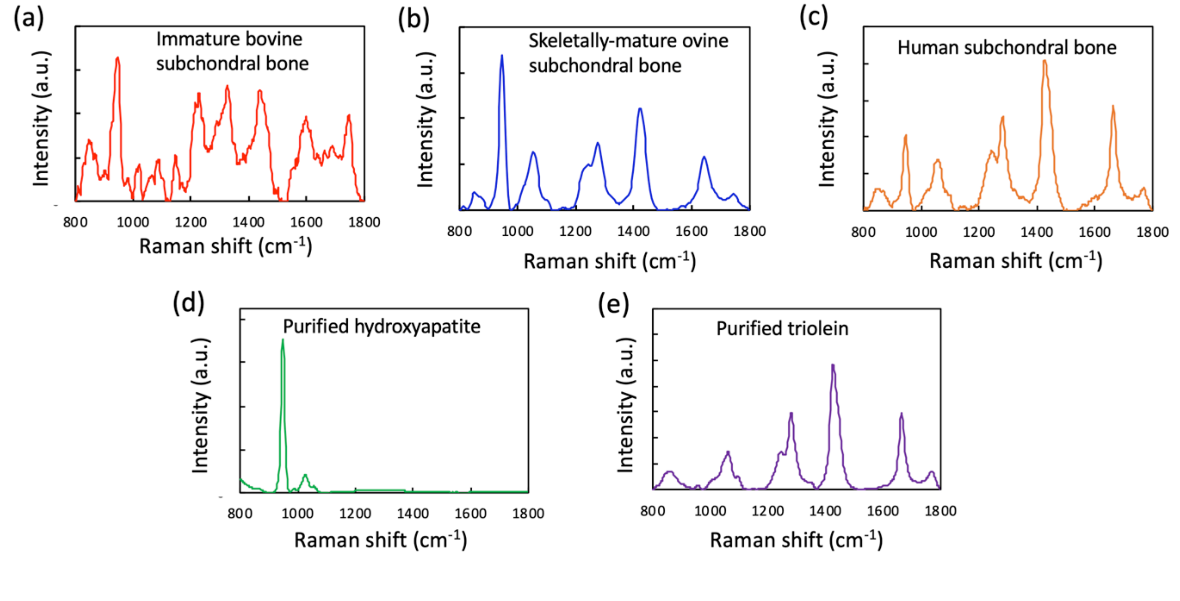


**Fig. S3. Raman spectra of subchondral bone** from (**a**) immature bovine, (**b**) skeletally mature ovine, and (**c**) human cartilage. These subchondral bone reference spectra are used as component spectra in regression analysis for bovine cartilage thickness measurements (Fig. 6), ovine cartilage analysis (Fig.1 and Fig. 7), and human cartilage analysis (Fig.4e). Spectra of purified chemical (**d**) hydroxyapatite powder and (**e**) triolein liquid.

***In situ* measurement assessments**

*In situ* intra-articular Raman needle diagnostics can encounter confounding factors in the synovial joint such as spectral interference from synovial fluid (SF), signal fluctuations from varying contact angles between the probe and articular cartilage surface, and sufficient acquisition time to acquire reliable Raman spectra. To examine the potential impact of these factors on Raman diagnostic measurements, a series of parametric studies were conducted on bovine cartilage explants, examining the effect of: 1) synovial fluid, 2) probe incidence angle, and 3) Raman integration time:

- Raman measurements were performed (using the ∅2mm fiber-optic ball lens) on immature bovine cartilage explants immersed in either a bath of PBS or SF (n=8 explants per group). Synovial fluid was procured from healthy mature bovine knee joints (Animal Technologies). Raman GAG scores were not significantly influenced by bathing fluid (Fig.S4a; p=0.92), suggesting that the synovial joint fluid environment does not impact Raman measurements.
- Raman measurements were performed with decreasing incidence angles between the probe tip and articular surface (90° [normal to surface], 80°, 70°, 60°, and 50°) (repeated measures on n=7 explants). A decrease of incidence angle to 70° had no significant effect on measured Raman GAG, COL, or $H_2O$ scores (Fig.S4b), indicating that a range of contact angles can be employed without compromising measurement accuracy.
- Raman measurements were performed on explants with varying spectral integration times (0.5, 1, 3, 10, 20, 30, 60, 90, and 120 seconds). There was no significant effect on integration time on Raman GAG, COL, and $H_2O$ scores (Fig.S4c).

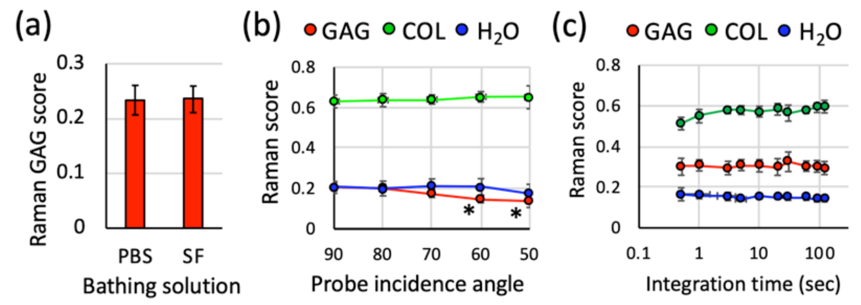


**Fig. S4.** (**a**) GAG score measurements of cartilage explants in a solution of PBS versus synovial fluid (SF). p=0.92 between groups. (**b**) GAG, COL, and $H_2O$ scores versus probe incidence angle. *p<0.05: indicates significant decrease below corresponding 90° angle Raman score value. (**c**) Effect of Raman acquisition (integration) time on GAG, COL, $H_2O$ scores of cartilage explants.

**Raman needle arthroscopy *in situ* synovial joint diagnostics**

*In situ* intra-articular Raman needle arthroscopy was performed on amputated intact bovine antebrachiocarpal joints (n=3). Measurements of cartilage ECM composition were performed before and after enzymatic cartilage degradation achieved via intra-articular injection of trypsin (10mL at 2000 μg/mL for 18 hours at 4°C). After Raman analysis, a full thickness chondral plug was explanted from each joint and subjected to DMMB assay to confirm enzymatic-induced GAG depletion. Cartilage GAG content was reduced: 4.3±0.6 %ww control vs 0.6±0.1 %ww trypsin-treated (Fig.S5a). Through a 10-gauge hypodermic needle trocar, *in situ* Raman probe measurements were executed with our ball lens (Fig.S5b). Similar to clinical joint arthroscopy, the joint was irrigated with saline prior to Raman measurements. The Raman probe was advanced through the syringe until it just contacted the articular joint surface and spectra measures were performed at the surface. Consistent with direct biochemical measurements, the derived Raman GAG score decreased with trypsin enzymatic GAG depletion, but COL and $H_2O$ scores were little changed (Fig.S5c).

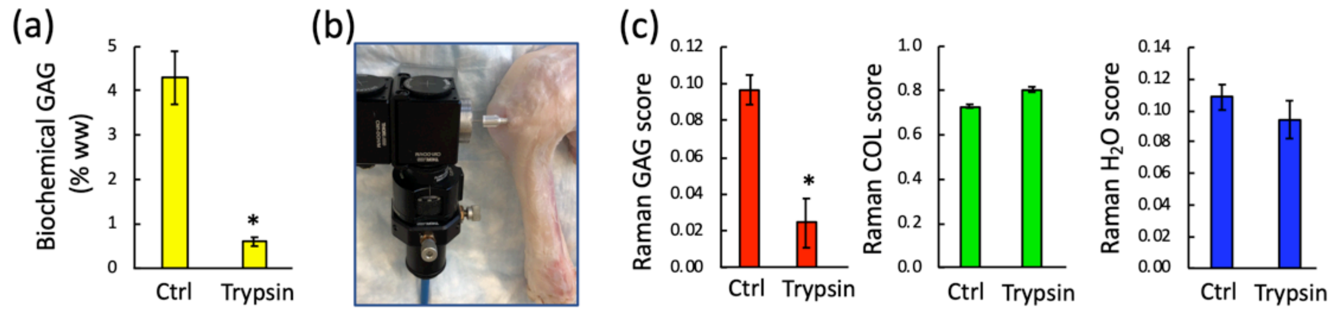


**Fig. S5**. **Raman arthroscopy *in situ* diagnostics**: (**a**) GAG reduction in intact bovine antebrachiocarpal joints after intra-articular trypsin treatment. (**b**) Intra-articular *in situ* diagnostics of an amputated bovine wrist joint with an intact synovial capsule. (**c**) Raman scores of GAG, COL, and $H_2O$ before (ctrl) and after enzymatic degenerative treatment (trypsin). *$p<0.01$: indicates a significant decrease below ctrl levels.

**Correlation parameters**

| Raman score (x) | Cartilage property (y) | Correlation function | $R^2$ value | p value |
|---|---|---|---|---|
| Raman GAG score | Biochemical GAG | y = 18.7x + 0.3 | 0.95 | < 0.001 |
| Raman COL score | Biochemical GAG | y = 22.1x + 18.5 | 0.91 | < 0.001 |
| Raman $H_2O$ score | Biochemical GAG | y = 19.4x + 5.8 | 0.03 | 0.24 |

**Table S1**: Correlation parameters of Raman scores versus biochemical GAG content in GAG depleted bovine cartilage explants (presented in Fig.2).

| Lens type | Raman score (x) | Cartilage property (y) | Correlation function | $R^2$ value | p value |
|---|---|---|---|---|---|
| Ball lens | Raman GAG score | Histology intensity | y = 1459x + 54 | 0.85 | < 0.001 |
| Ball lens | Raman COL score | Histology intensity | y = 1896x + 1583 | 0.77 | < 0.001 |
| Ball lens | Raman $H_2O$ score | Histology intensity | y = 2263x + 730 | 0.24 | 0.05 |
| Convex lens | Raman GAG score | Histology intensity | y = 4786x + 135 | 0.40 | < 0.01 |
| Convex lens | Raman COL score | Histology intensity | y = 4329x + 3672 | 0.26 | 0.04 |
| Convex lens | Raman $H_2O$ score | Histology intensity | y = 7790x + 2163 | 0.14 | 0.13 |

**Table S2**: Correlation parameters of Raman scores versus Safranin-O histology intensity from trypsin-treated cartilage explants (presented in Fig.3).

| Lens type | Raman score (x) | Cartilage property (y) | Correlation function | $R^2$ value | p value |
|---|---|---|---|---|---|
| Ball lens | Raman GAG score | Biochemical GAG | y = 48.2x + 0.6 | 0.66 | < 0.001 |
| Ball lens | Raman COL score | Biochemical GAG | y = -31.9x + 28.5 | 0.50 | < 0.01 |
| Ball lens | Raman $H_2O$ score | Biochemical GAG | y = 23.8x + 1.2 | 0.13 | 0.23 |
| Ball lens | Raman GAG score | SOFG uptake score | y = 39.4x + 3.3 | 0.58 | < 0.01 |
| Convex lens | Raman GAG score | Biochemical GAG | y = 52.7x - 2.4 | 0.41 | 0.02 |
| Convex lens | Raman COL score | Biochemical GAG | y = -39.5x + 32.5 | 0.22 | 0.11 |
| Convex lens | Raman $H_2O$ score | Biochemical GAG | y = 24.3x - 0.6 | 0.02 | 0.65 |
| Convex lens | Raman GAG score | SOFG uptake score | y = 51.3x + 6.6 | 0.53 | < 0.01 |

**Table S3**: Correlation parameters of Raman scores versus biochemical GAG content and SOFG stain uptake scores in human cartilage explants (presented in Fig.4).

| Cartilage property (x) | Raman score (y) | Correlation function | $R^2$ value | p value |
|---|---|---|---|---|
| Cartilage thickness | Raman bone score | $y = 1.0e^{-0.91x}$ | 0.90 | < 0.001 |
| Cartilage thickness | Raman GAG score | y = 0.11x + 0.05 | 0.85 | < 0.001 |
| Cartilage thickness | Raman $H_2O$ score | y = 0.05x | 0.91 | < 0.001 |
| Cartilage thickness | Raman COL score | y = 0.20x + 0.14 | 0.79 | < 0.001 |

**Table S4**: Correlation parameters of Raman scores versus cartilage thickness (presented in Fig.6).

**Statistical analysis**

An unpaired Student's t-tests (two-tailed) was used to compare Raman GAG scores of explants between PBS and SF (Fig.S4a). One-way analysis of variances (ANOVAs) were performed ($\alpha$=0.05) to determine the effect of incidence angle on Raman GAG score or Raman $H_2O$ score. A paired Student's t-test was used to compare GAG content and Raman GAG scores before and after trypsin treatment (Fig.S5).
